# A prospective, observational study on conversion of Clinically Isolated Syndrome to Multiple Sclerosis during 4-year period (MS NEO study) in Taiwan

**DOI:** 10.1101/385898

**Authors:** Long-Sun Ro, Chih-Chao Yang, Rong-Kuo Lyu, Kon-Ping Lin, Tzung-Chang Tsai, Shiang-Ru Lu, Kuo-Hsuan Chang, Li-Chieh Huang, Ching-Piao Tsai

**Affiliations:** Department of Neurology, Chang Gung Memorial Hospital-Linkou Medical Center, Chang Gung University College of Medicine, Taoyuan, Taiwan (R.O.C.); Division of Nephrology, Department of Internal Medicine, Department of Internal Medicine, Kaohsiung Chang Gung Memorial Hospital and Chang Gung University College of Medicine, Kaohsiung, Taiwan (R.O.C.); Department of Neurology, Neurological Institute, Taipei Veterans General Hospital and National Yang-Ming University, Taipei, Taiwan (R.O.C.); Department of Neurology, Taichung Hospital, Ministry of Health and Welfare, Taichung, Taiwan (R.O.C.); Department of Neurology, Faculty of Medicine, College of Medicine, Kaohsiung Medical University, Taiwan; Department of Neurology, Kaohsiung Medical University Hospital, Taiwan. (R.O.C.); Department of Neurology, Chang Gung Memorial Hospital-Linkou Medical Center, Chang Gung University College of Medicine, Taipei, Taiwan (R.O.C.); Merck Biopharma (Taiwan), Taipei, Taiwan (R.O.C.); Department of Biotechnology, Asia Unisersity, Taichung, Taiwan. Taipei Beito Health Management Hospital, Taipei, Taiwan

## Abstract

**Importance:** CIS to MS conversion rates vary depending on population cohorts, initial manifestations, and durations of follow-up.

**Objective:** To investigate conversion rate of patients from CIS to MS and the prognostic significance of demographic and clinical variables in Taiwanese population.

**Design:** Nationwide, prospective, multi-centric, observational study from November 2008 to November 2014 with 4 years follow-up.

**Setting:** Multi-centre setting at 5 institutions in Taiwan.

**Participants:** 152 patients having single clinical event potentially suggestive of MS in last 2 years were enrolled as consecutive sample. 33 patients were lost to follow-up and 16 patients did not complete the study.103 patients completed the study.

**Intervention(s) (for clinical trials) or Exposure(s) (for observational studies):** Natural progression from first episode of CIS to MS or NMO was observed.

**Main Outcome(s) and Measure(s):** Variables analysed were ‘proportion of patients converting to MS or NMO after first episode of CIS’, ‘duration between first episode of neurological event and diagnosis of MS’, ‘status of anti-AQP4 IgG’ and ‘length of longest contiguous spinal cord lesion in MS patients’. Association between baseline characteristics and progression to MS from CIS was analyzed using multiple logistic regression. Multivariate time dependent effect of baseline characteristics on progression to MS was plotted.

**Results:** 14.5% patients with CIS converted to MS after 1.1 ± 1.0 years with greater predisposition (18.8%) in those having syndromes referable to the cerebral hemispheres. Conversion rate from ON to MS was 9.7%. 90.9% patients had benign disease course. 46.7% patients had abnormal MRIs at baseline, with 0.6±0.5 contrast enhanced lesions. ‘Below normal BMI’ and ‘MRI lesion load (≥ 4 lesions)’ were identified as risk indicators for the development of MS. Only 4.5% were positive for anti-AQP4 antibody in MS patients and amongst them, 80% were NMO patients as diagnosed by modern criteria.

**Conclusions and Relevance:** ‘Below normal BMI’ and ‘number of demyelinating lesions (≥4)’ are significant predictors of conversion from CIS to MS. A low conversion rate to MS in Taiwanese CIS patients and majority of them having a benign course and minimal disability suggest the roles of geographic, genetic and ethnic factors.

**Trial Registration:** Non-trial observational study.

## Introduction

CIS is the first clinical demyelinating event lasting ≥24 hours, prodromal to MS. It is also a stage at which intervention could be effective in reducing chances of conversion to MS. Early risk stratification in CIS helps to counsel the patients about prognosis and to expedite treatment initiation with the goal of reducing future morbidity [1,2]. Clinical phenotype of CIS includes spinal cord syndrome (50%), optic neuritis (25%), brain stem and/or cerebellar syndrome (15%), or occasionally cerebral hemispheric dysfunction [3,4].

Neuromyelitis optica (NMO) and acute demyelinating encephalomyelitis are the important differential diagnosis of CIS [4]. Many patients with NMO have detectable serum IgG antibodies against the water channel aquaporin-4 (AQP4–immunoglobulin G [IgG]) which are very specific for clinically diagnosed NMO. Currently, routine testing for AQP4-IgG antibodies is considered in high-risk individuals due to similar initial disease course in NMO and MS [5]. MS is chiefly diagnosed by clinical acumen supported by investigations, including T2-weighted magnetic resonance imaging (MRI), CSF evaluation, and visual evoked potential [5].

Longitudinally-extensive spinal cord lesions are uncommon and are suggestive of NMO[4]. MRI remains the most important surrogate marker for predicting the risk of a second event in CIS patients (i.e conversion to MS) but the clinical outcome remains unpredictable due to the high variability of this disease among individuals. Thus, a need remains for auxiliary markers that could provide additional information about the disease course.

Various prospective, longitudinal cohort studies in CIS patients have reported the rates of conversion to MS. The conversion rate from CIS to MS is reported to lie between 30%–82% by different global studies [4,6,7,8,9]. The variability in conversion rates is likely attributable to the cohort studied, geographic area, ethnic group, initial manifestation, and the duration of follow-up. Moreover, factors like difference in patient selection method, diagnostic criteria, and study design also cause variability.

Numerous factors influence the conversion to MS after the first episode of CIS. Of the epidemiological risk factors, female gender and young age group are shown to be predictive of MS after a CIS [6–11,22]. Long term studies have shown that number and activity of MRI lesions and spinal cord abnormalities at the time of presentation are the strongest predictor of conversion to MS [2,12]. As regards the location, Barkhof *et al* has suggested that periventricular demyelinating lesions foresee conversion to MS in patients with CIS [13]. A shorter interval to a second episode and the number of relapses in the first 2 years are also indicators of a poor prognosis [14].

In Taiwan, a retrospective single centre study revealed that conversion rate to MS was 14.3% in patients with optic neuritis. Female gender, retro bulbar type ON, MRI abnormalities, elevated CSF IgG index, and recurrent attacks were identified as risk factors [6]. Another study reported 9 year cumulative incidence rate of MS in patients with ON to be 1.02% with a predominance in females and younger patients.

None of the studies has prospectively analysed the conversion of CIS to MS in Taiwanese population. The precise likelihood of further neurological events and conversion to MS is essential both for patients and neurologists pertaining to timely initiation of disease modifying therapies.

The current study was planned with the primary objective to evaluate the conversion rate from CIS to MS over a 4-year period in Taiwanese patients. The secondary objectives were to assess the relationship between CIS and MS, the status of anti-AQP4 IgG in MS patients, to determine the length of longest contiguous spinal cord lesion in MS patients and to evaluate association between baseline demographic/ disease characteristics with conversion to MS.

## Materials and methods

### Study Design

The MS Neo Study is a nationwide, multi-centre, prospective, observational study. Briefly, between November 2008 and November 2014, 152 patients who had single clinical event potentially suggestive of MS within the last 2 years were enrolled at 5 institutions in Taiwan.

The study was approved by the institutional review board and ethics committee of each institute and performed in accordance with the principles of the Declaration of Helsinki. Written informed consent was obtained from all patients.

The study participants were encouraged to follow up at 12 – 16 weeks in case of no event, and in case of a relapse until diagnosis of MS or other disease. A telephonic interview was conducted at quarterly intervals. 33 patients were lost to follow-up and 16 patients did not complete the study due to other reasons. Subsequently, 103 patients completed the study. The study participants had a minimum life expectancy of ≥12 weeks.

### Data collection

During the baseline and the follow up visits, the neurological disability was clinically assessed by the Expanded Disability Status Scale (EDSS). In all patients, baseline and follow-up MRI scans were performed. MRI study included axial and sagittal images of the brain and spinal cord obtained by T1, T2, FLAIR and T1 post-contrast sequences.

The AQP4 antibody titre was determined at the baseline visit, follow up visits and at the completion of the study. At the time of study completion, the type of diagnosed disease was determined along with ophthalmological and neurological examination. The date of MS conversion was noted. The adverse events (AE) were assessed at each of the follow up visits.

### Study Endpoint

MS was defined by the McDonald criteria 2005 and compared with McDonald criteria 2010 after study completion. Variables analysed were ‘proportion of patients converting to MS or NMO after first episode of CIS’, ‘duration between first episode of neurological event and diagnosis of MS’, ‘status of anti-AQP4 IgG’ and ‘length of longest contiguous spinal cord lesion in MS patients’. Association between baseline characteristics and progression to MS from CIS was analyzed using multiple logistic regression. Multivariate time dependent effect of baseline characteristics on progression to MS was plotted.

### Statistical considerations

All patients enrolled were included in the analysis. The primary endpoint was summarized as qualitative variable and presented with number of subjects, and percentage. Multiple logistic regression was used to determine the association between baseline demographics/disease characteristics and progression to MS from CIS. A probability of p ≤0.05 was accepted as significant. Kaplan-Meier graph was plotted using duration between initial CIS to MS diagnosis. A forest plot of multivariate time dependent effect of baseline characteristics on the progression to MS was plotted. All analyses were performed using Statistical Analysis Software(SAS) version 9.3.

## Results

### Epidemiological and clinical data

A total of 152 CIS patients were included in the study and analysed. These patients had presented to neurology services between 4^th^November 2008 and 20^th^ November 2014. The flowchart of patient selection is depicted in Table 1. All patients enrolled were included in the statistical analysis.

**Table 1:**
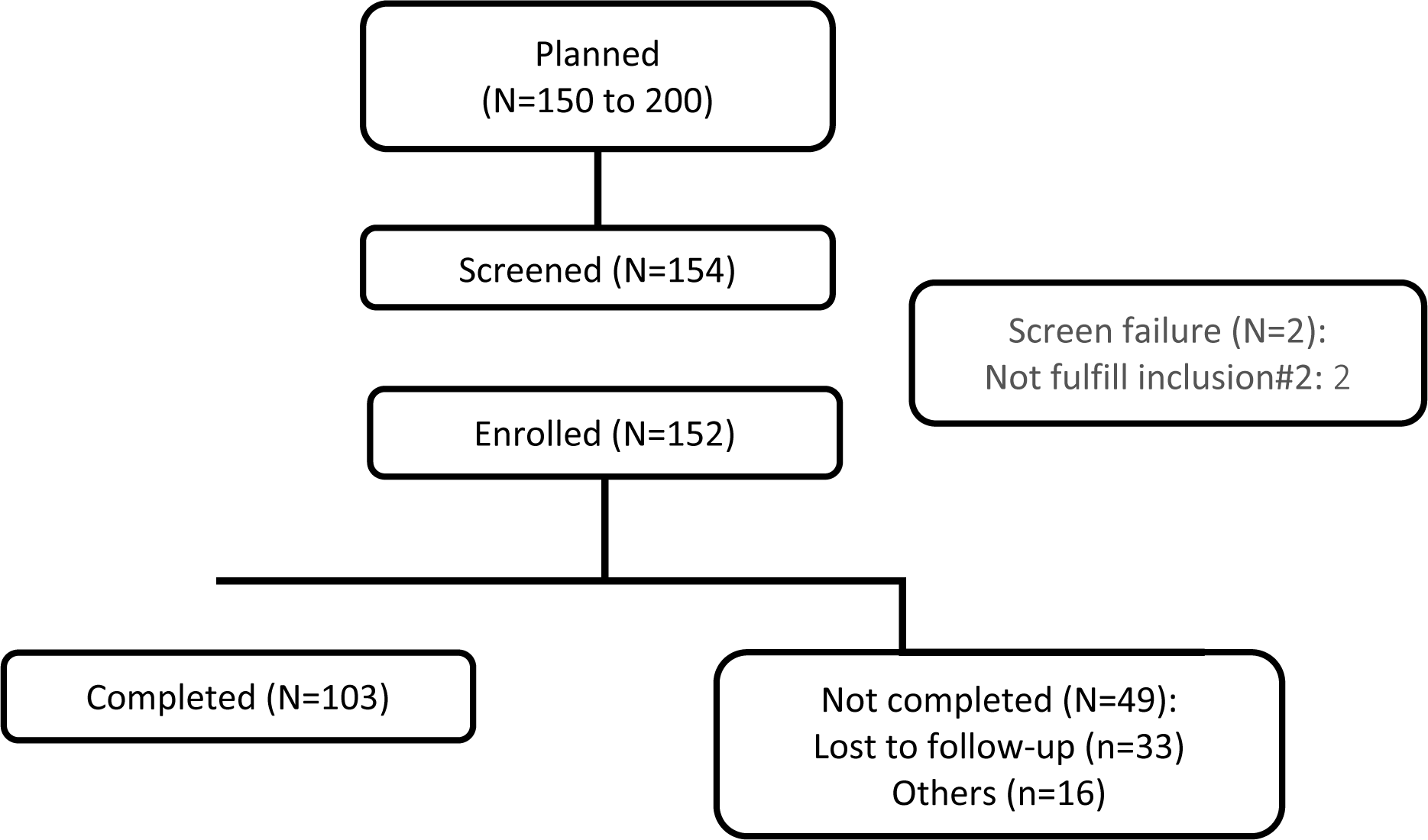
Consort diagram of patient flow

Among the 152 CIS patients included in the study, 116 (76.3%) were women and the mean age at baseline was 38.6 ± 13.3 years (interquartile range, 27.2–50.2). The demographic characteristics of CIS patients at the baseline are shown in Table 2.

**Table 2:** Baseline characteristics of CIS patients

| Parameter | Statistics | Results |
| --- | --- | --- |
| Gender | N | 152 |
|  | Female | 116 (76.3%) |
|  | Males | 36 (23.7%) |
| Race | N | 152 |
|  | Asians | 152 (100%) |
| Age (Year) | N | 152 |
|  | Mean (STD) | 38.6 (13.3) |
|  | Median | 38.5 |
|  | (Q1 ,Q3) | (27.2, 50.2) |
|  | (Min, Max) | (9.4, 67.3) |
| Height (cm) | N | 151 |
|  | Mean (STD) | 160.3 (7.5) |
|  | Median | 160.0 |
|  | (Q1 ,Q3) | (155.0, 165.0) |
|  | (Min, Max) | (139.0, 180.0) |
| Weight (kg) | N | 150 |
|  | Mean (STD) | 58.2 (11.4) |
|  | Median | 57.0 |
|  | (Q1 ,Q3) | (50.3, 64.5) |
|  | (Min, Max) | (33.0, 124.0) |
| BMI (kg/m<sup>2</sup>) | N | 150 |
|  | Mean (STD) | 22.6 (3.7) |
|  | Median | 22.3 |
|  | (Q1 ,Q3) | (19.9, 24.8) |
|  | (Min, Max) | (15.1, 40.9) |
| BMI: Body Mass Index. Subject Didn't Collect Weight: 06-012, Subject Didn't Collect Weight and Height: 01-027 |  |  |

We analysed the association between baseline demographic characteristics (age, gender and BMI) in CIS patients and the conversion rate to MS using multiple logistic regression and did not find any statistically significant association as shown in Table 3. Association between race and MS conversion could not be analyzed as all the enrolled subjects were Asians.

**Table 3:** Association between baseline demographics/disease characteristics and conversion to MS

| Parameter | Category | Coefficient | E<sup>^</sup>Coefficient (OR) | P-value |
| --- | --- | --- | --- | --- |
| Age (Year) |  | −0.016 | 0.984 | 0.363 |
| BMI (kg/m<sup>2</sup>) |  | 0.017 | 1.018 | 0.778 |
| Gender | Male vs. Female | −0.180 | 0.835 | 0.757 |
| Location of Longest Lesion | Peripheral vs. Central | −1.131 | 0.323 | 0.459 |
|  | Holospine vs. Central | −13.551 | <0.001 | 0.974 |
|  | Central+Peripheral vs. Central | 13.683 | >999.999 | 0.990 |
|  | NA vs. Central | −2.293 | 0.101 | 0.014 |
| EDSS Score |  | 0.170 | 1.186 | 0.135 |
| Analyzed using multiple logistic regression to evaluate association between baseline demographics/disease characteristics and conversion to MS from CIS.<br>BMI: Body Mass Index; EDSS: Expanded Disability Status Scale |  |  |  |  |

Of these patients, 142 (93.4%) presented with CIS at the baseline visit. The type of episode was optic neuritis (ON) in 72(47.4%) patients and transverse myelitis (TM) in 47(30.9%) patients. The clinical episode in one patient (0.7%) consisted of simultaneous optic neuritis and transverse myelitis. Apart from ON and TM, 32(21.1%) patients experienced other syndromes such as encephalopathy, brainstem encephalitis, right hemispheric attack, and acute disseminated encephalomyelitis (ADEM).

According to the established diagnostic criteria for MS^2^, the conversion rate to MS was 14.5% in our study, with an average duration of 1.1 ± 1.0 years (interquartile range, 0.2–1.9). 5 (3.3%) patients developed NMO. Thus conversion rate to collectively either MS or NMO was 17.8% during 4-year observation period. MS developed in 7 out of 72 (9.7%) who initially presented with ON and in 8 out of 47 (17.0%) with a spinal cord syndrome. Six of 32 patients (18.8%) with other syndromes converted to MS and one patient who had presented with simultaneous event of ON and TM also converted to MS.

Eleven patients (50%) developed MS within six months of first clinical isolated syndrome and three (13.6%) converted to MS within a year. The same finding has been depicted in the Kaplan-Meier plot of duration between initial CIS to MS diagnosis. (Fig 1)

**Figure 1:**
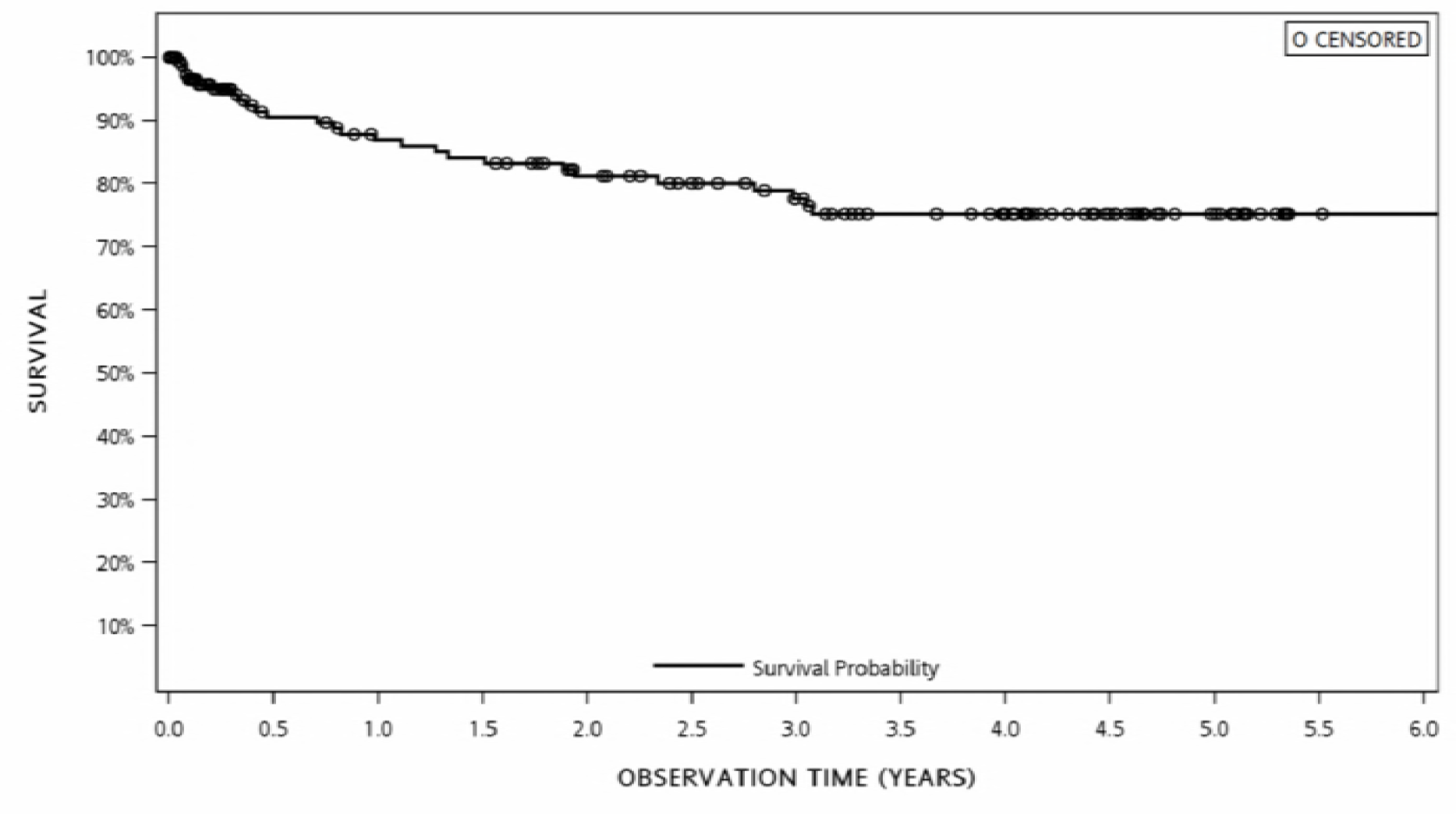
KM plot of duration between initial CIS to MS diagnosis.

The mean baseline EDSS score of patients who developed multiple sclerosis was 2.4 ± 2.0 (range 0.0 to 9.0). The score was found to be between 0.0 to 4.5 in 20 (90.9%) patients with a benign disease course and score was between 5.0 to 10.0 in 2 (9.1%) subjects indicating that assistance was required for walking. This extent of neurological disability increased gradually with a score of 3.8 ± 2.6 at the time of relapse. The mean EDSS score in NMO subjects was 4.2 ± 2.3 at baseline. There was no significant association between the EDSS score at baseline and conversion to MS (P = 0.135). [Tables 3 and 4]

**Table 4:**
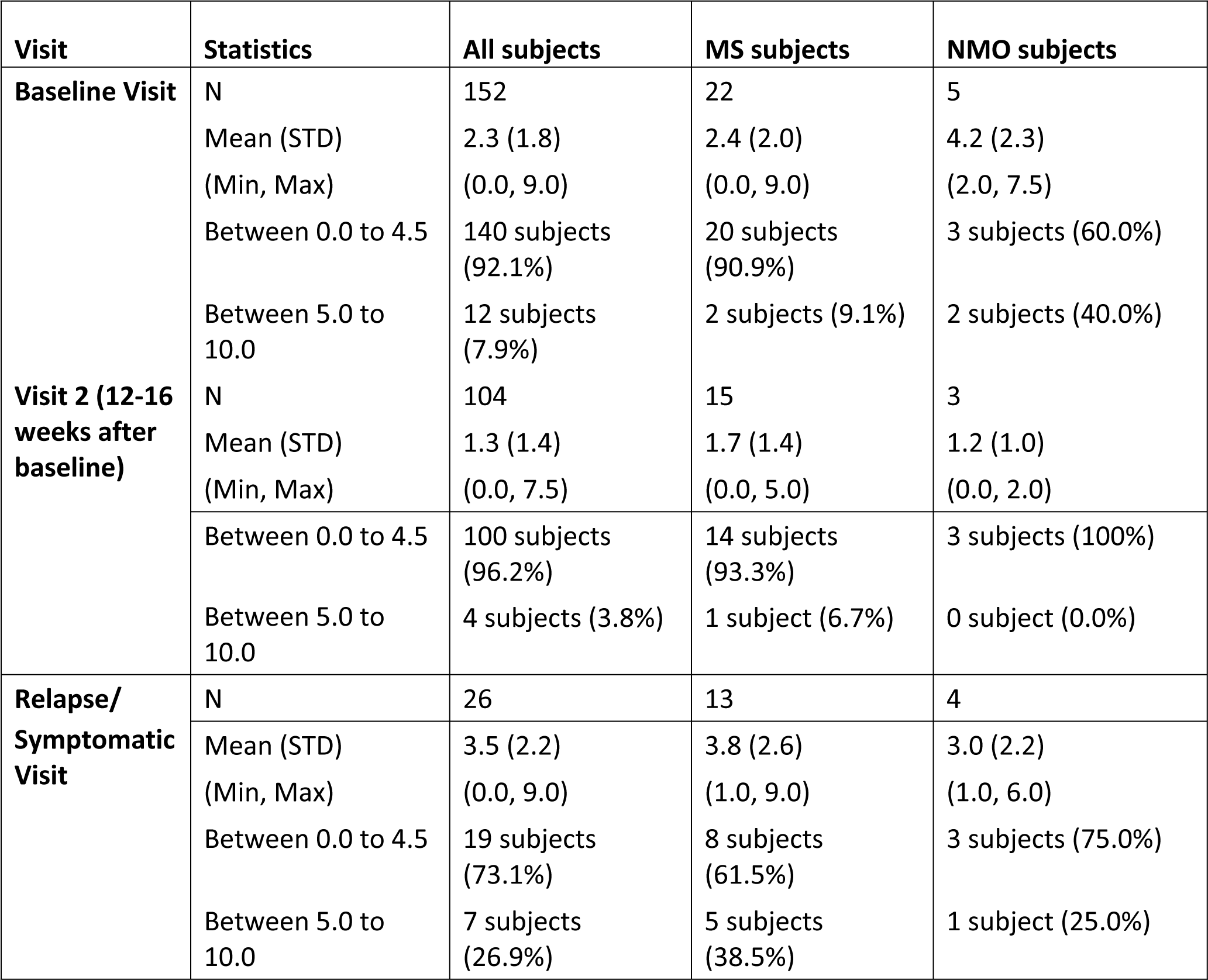
EDSS score at baseline and follow-up visits

### Neuroimaging study

Seven (46.7%) patients with an initial episode of optic neuritis who converted to MS were having an abnormal brain MRI scan at the baseline. The average number of contrast enhanced lesions in these patients was 0.6 ± 0.5 (interquartile range, 0.0 – 1.0) and only one patient fulfilled the Barkhoff criteria in the baseline brain MRI.

The spinal MRI revealed that the average number of lesions were 1.2 ± 1.1. The length of the longest spinal cord lesion at baseline was 2.0 ± 0.8 (range 1.0 – 3.0) in patients who converted to MS and 3.5 ± 0.7 (range 3.0 – 4.0) in patients who developed NMO. The location of the longest lesion was central in 50% of patients followed by peripheral (25%) and holospine (16.7%). Multiple logistic regression revealed no significant association between the location of the longest lesion in baseline MRI and conversion to MS. (Table 3)

### Anti-AQP4 IgG antibody status

Only one patient (4.5%) of all those who developed MS was with positive anti-AQP4 antibody and this patient was tested positive during the relapse/symptomatic visit. As regards the patients who developed NMO, 4 of 5 (80.0%) were positive for anti-AQP4 antibody.

A multivariate analysis was conducted incorporating time-dependent effect of demographic and disease characteristics, clinical and radiological data available at baseline. In the multivariate analysis (Fig 2), our study cohort revealed that ‘age’, ‘gender’ and ‘EDSS’ had non-significant association with MS conversion rate but a trend was noted. ‘BMI’ and ‘number of lesions’ had significant association with probability of MS conversion.

**Figure 2:**
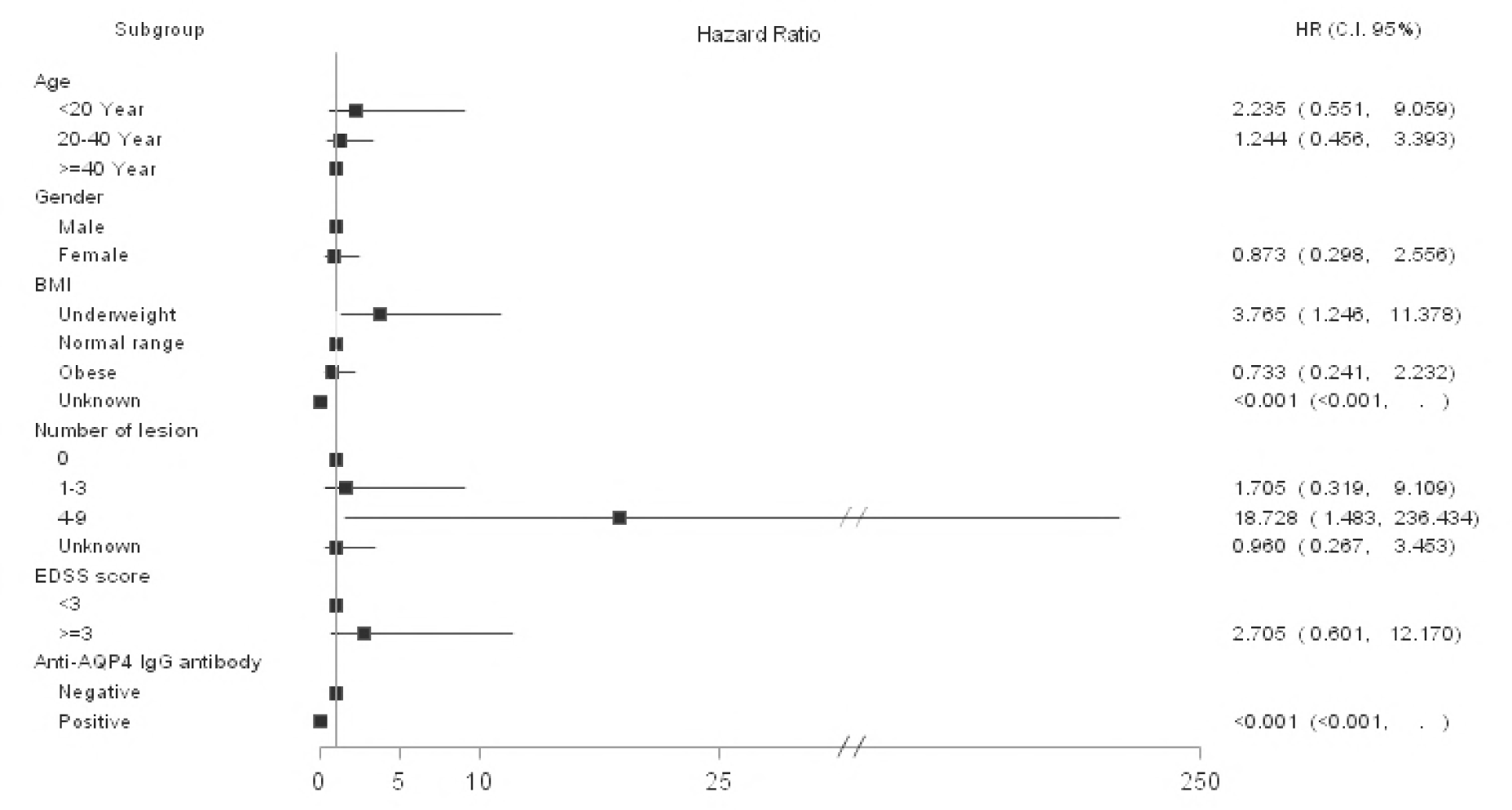
Forest Plot of Multivariate Time Dependent Effect of Baseline Demographic/ Disease Characteristics on the Conversion to MS (Enrolled population)

### Adverse events

Of the 152 subjects enrolled in the study, 98 (64.5%) patients had reported at least one AE. Eleven (7.2%) patients reported with at least one severe AE. The most common AEs reported were anxiety and tear film insufficiency in 8(5.3%) patients. Cough, dry eye, and headache was complained of in 7(4.6%) patients while 6(3.9%) patients had allergic conjunctivitis. Chronic conjunctivitis, constipation and hypertension was seen in 5 (3.3%) patients. A total of 6 (3.9%) patients reported at least one serious adverse event (SAE).

## Discussion

The present study provides prospectively acquired epidemiological, clinical and MRI data over a period averaging 4 years from CIS onset in Taiwanese population. The overall conversion rate to MS was 14.5% in patients presenting with CIS. The conversion rates to MS in our study are considerably lower than those observed in most of the trials including the optic neuritis treatment trial. (Table 5)

**Table 5:** Previous studies of the conversion rate of multiple sclerosis following idiopathic optic neuritis

| **Study** | **Year** | **No. of patients (n)** | **Conversion rate** | **Follow up period** |
| --- | --- | --- | --- | --- |
| Optic neuritis study group (USA)<sup>7</sup> | 1997 | 389 | 50% | 15 years |
| Fisniku et al (UK)<sup>12</sup> | 2008 | 54 | 65% | 20 years |
| Kuhle et al (UK) <sup>8</sup> | 2015 | 1027 | 59.5% | 4.31 years |
| ZhonghuaNeiKe (China)<sup>16</sup> | 2016 | 151 | 30.5% | 44.11±17.62 months |
| Saxena et al<sup>17</sup> | 2016 | 83 | 28% | 18 months |
| Marques et al<sup>18</sup> | 2014 | 42 | 23.8% | 5 year |

In Japan, a statistical survey of ON revealed that rate of conversion of ON to MS was 8.3% [15]. A partially retrospective and longitudinal study in Mexico showed that the percentage of patients that converted from primary idiopathic ON to a clinically definite MS in a follow-up period of 7 years was 12% [19]. The conversion rates of ON to MS in our study were similar to these studies (9.7%). This implies an important role of geographic, genetic and ethnic factors in conversion of CIS, in particular ON to MS. Moreover, the finding of overall low conversion rate to MS of Taiwanese patients complement the results of the earlier retrospective studies which reveals low risk of MS in Taiwan [6,9].

Though some studies have shown that optic neuritis is associated with a lower risk of MS, the risk is comparable across different CIS subtypes[4]. In this report, CIS patients presenting with other syndromes had greater predisposition to convert than the patients with ON and/or TM at the time of presentation. These findings may be attributable to differences in patient inclusion criteria, diagnostic criteria for MS and the duration of the follow-up period in various studies.

Among the patients who converted to MS, a female preponderance was seen in the ratio of 3:1. This finding is consistent with the well described gender predilection of MS [1,4,6,7,20]. The mean age at the onset of neurological symptoms (38.6 years) in our study is similar to the mean age reported by the ONTT and other major trials [7,8,12,17,18,20]. However, contrary to other studies, we did not find any association between mean age at onset and gender with the risk to MS conversion [7,22].

Our study reported that half of the patients developed MS within six months of first CIS and the rest within a year. Analogous to this study, Kuhle *et al*, in a large international cohort, found the duration to conversion was in the range of 7 months to 2 years [8]. Marques *et al* reported that majority of ON patients converted to MS during the first follow-up year [18].

The duration to convert to MS from ON is generally associated with the number and location of demyelinating lesions of the brain and spinal cord [6,22]. In this study, there was a clear distinction in the conversion rates between patients with and without evidence of prior demyelination on brain MRI. In keeping with several previous studies, we confirmed the fact that abnormal baseline brain MRI is a strong predictor for the development of MS in idiopathic ON patients [7,8,12].

The multiple sclerosis group in our study exhibited a narrow spectrum of extent of disability. Whilst 91% had a benign course with minimal disability (EDSS < 4.5), only a few patients (9%) developed secondary progressive MS with increased neurological disability (EDSS 5 – 10). The percentage of people with benign multiple sclerosis in our cohort was higher than reported on other cohort of patients. In a study with a follow up period of 20 years, 42% of patients had substantial neurologic disability with a median EDSS score of 6.5 [12]. Nonetheless, natural history of the disease suggests that the number of patients with a benign relapsing remitting MS reduces steadily with increasing duration of follow up.

In our study, 80% patients who developed NMO were tested positive for anti-AQP4 antibody and 4.7% patients fulfilled the criteria for NMOSD suggestive of a high risk of future relapses [5].

A multivariate analysis was conducted incorporating time dependent effect of demographic and disease characteristics on the conversion to MS. (fig 2) Greater risk of relapse in younger patients in the age group of ˂ 20 years compared to patients aged ≥ 40 years at disease onset, however, the statistical significance of this result could not be demonstrated. Gender does not seem to be a risk factor for conversion to MS or a further relapse, in line with the gender predilection stated in literature[23,24]. Nonetheless, we cannot exclude the probability that a longer follow-up period might demonstrate a difference in MS conversion among genders. Another study by Tintore M et al. [25] reported lack of gender predilection for conversion rate from CIS to MS in multivariate analysis with an observation period of 18 years.

Our study revealed significant association between the baseline BMI and conversion to MS. It showed that being underweight is a significant risk factor for conversion from CIS to MS.

The analysis confirms that MRI at baseline is a robust factor predictive of conversion to multiple sclerosis. Almost half of the patients who converted to MS after 4 years had demyelinating lesions of the brain and spinal cord on MRI scan at baseline with Gd contrast enhanced lesions. The number of lesions ranging from 4 to 9 on MRI scan at baseline is a significant risk factor for conversion to MS which is consistent with the findings by Tintore M et al. [25] Another important clinical factor related to multiple sclerosis is the extent of neurological disability assessed by EDSS score. Our study showed that patients with an EDSS score ≥ 3 display a higher risk of conversion to MS. It was observed that presence of anti-AQP4 IgG antibody indicates less likely possibility of conversion of CIS to MS. Patients who tested positive for anti-AQP4 antibody had NMO as diagnosed by modern criteria.

Many CIS episodes are mild and resolve without therapeutic intervention. Currently available treatment have unequivocal efficiency and may slow the conversion to RRMS, but could be avoided for patients with benign disease course [21]. During an average 4 year follow up period, 7.2% patients reported at least one adverse event, most common being anxiety and tear film insufficiency. Six patients suffered from serious adverse event. Thus, with the introduction of highly effective disease-modifying treatments for MS, there is a need for risk assessment to aptly select therapies for these patients due to the associated serious adverse effects.

A key strength of our study was inclusion of patients from majority of medical centres in the country. However, the study had some limitations, including a relatively small sample size. We could not assess the vitamin D levels in this study. Also, we could not determine the effect of ethnicity on the clinical course as our study included only Asian population. Furthermore, the variables assessed were not adjusted for ancestry and genetic variants with an inherent MS risk, thereby limiting the generalization of our results. EDSS scale used has inherent limitations like intra-observer variability and heavy weightage given to physical disability with limited assessment of key clinical features. Furthermore, MS conversion rate may have been underestimated as only clinical MS conversion was considered and not the new silent subclinical lesions.

## Acknowledgments

The authors would like to acknowledge the medical writing support provided by Dr. Pratishtha Banga, MD from WriterMD Medical Writing Consultancy

## References

1. Thouvenot É. Update on clinically isolated syndrome. Presse Med. 2015; 44(4 Pt 2):e121–36.

2. Milo R, Miller A. Revised diagnostic criteria of multiple sclerosis. Autoimmun Rev. 2014; 13(4-5):518–24.

3. Eriksson M, Andersen O, Runmarker B. Long-term follow-up of patients with clinically isolated syndromes, relapsing-remitting and secondary progressive multiple sclerosis. MultScler 2003; 9:260–74.

4. O’Riordan JI, Thompson AJ, Kingsley DP, MacManus DG, Kendall BE, Rudge P et al. The prognostic value of brain MRI in clinically isolated syndromes of the CNS. A 10-year follow-up. Brain. 1998 Mar; 121(Pt 3):495–503.

5. Wingerchuk DM, Banwell B, Bennett JL, Cabre P, Carroll W, Chitnis T, et al. International Panel for NMO Diagnosis.. International consensus diagnostic criteria for neuromyelitis optica spectrum disorders. Neurology. 2015 14; 85(2):177–89.

6. Lin YC, Yen MY, Hsu WM, Lee HC, Wang AG. Low conversion rate to multiple sclerosis in idiopathic optic neuritis patients in Taiwan. Jpn J Ophthalmol. 2006;50(2):170–5.

7. Optic Neuritis Study Group. Multiple sclerosis risk after optic neuritis: final optic neuritis treatment trial follow-up. Arch Neurol 2008;65:727–32

8. Kuhle J, Disanto G, Dobson R, Adiutori R, Bianchi L, Topping J, et al. Conversion from clinically isolated syndrome to multiple sclerosis: A large multicentre study. MultScler. 2015 Jul;21(8):1013–24.

9. Woung LC, Peng PH, Liu CC, Tsai CY, Wang KC, Lee WJ, et al. A nine-year population-based cohort study on the risk of multiple sclerosis in patients with optic neuritis. Tohoku J Exp Med. 2013; 231(3):171–7.

10. Dobson R, Ramagopalan S, Giovannoni G. The effect of gender in clinically isolated syndrome (CIS): a meta-analysis. MultScler 2012;18:600–4

11. Mowry EM, Pesic M, Grimes B, Deen SR, Bacchetti P, Waubant E. Clinical predictors of early second event in patients with clinically isolated syndrome. J Neurol 2009;256: 1061–6.

12. Fisniku LK, Brex PA, Altmann DR, Miszkiel KA, Benton CE, Lanyon R, et al. Disability and T2 MRI lesions: a 20-year follow-up of patientswith relapse onset of multiple sclerosis. Brain. 2008 Mar;131(Pt 3):808–17.

13. Barkhof F, Filippi M, Miller DH, et al. Comparison of MRI criteria at first presentation to predict conversion to clinically definite multiple sclerosis. Brain 1997; 120:2059–69

14. Scalfari A, Neuhaus A, Daumer M, et al. Onset of secondary progressive phase and longterm evolution of multiple sclerosis. J NeurolNeurosurg Psych 2014;85:67–75

15. Isayama Y, Takahashi T, Shimoyoma T, et al. Acute optic neuritis and multiple sclerosis. Neurology 1982; 32:73–76

16. Bi CF, Qian HR, Peng LJ, Mao LL, Huang X, Xia DY et al. The Correlation Factor Analysis for Conversion of Clinically Isolated Syndrome to Multiple Sclerosis and Neuromyelitis Optica. Zhonghua Nei Ke Za Zhi 55 (6), 460–465. 6 2016.

17. Saxena R, Phuljhele S, Menon V, Gadaginamath S, Sinha A, Sharma P. Clinical profile and short-term outcomes of optic neuritis patients in India. Ind J of Ophthal. 2014; 62(3):265–267.

18. Marques IB, Matias F, Silva ED, Cunha L, Sousa L. Risk of multiple sclerosis after optic neuritis in patients with normal baseline brain MRI. J Clin Neurosci. 2014; 21(4):583–6.

19. Corona-Vazquez T, Ruiz-Scadoval J, Arriada-Mendicoa N. Optic neuritis progression to multiple sclerosis. ActaNeurolScand 1997;95:85–89

20. Chang YC, Wu WC, Tsai RK. Prognosis of Taiwanese patients with isolated optic neuritis after intravenous methylprednisolone pulse therapy. J Formos Med Assoc. 2007;106(8):656–63.

21. Miller DH, Chard DT, Ciccarelli O. Clinically isolated syndromes. Lancet Neurol. 2012;11(2):157–69.

22. R. Alroughani, J. Al Hashel, S. Lamdhade, and S. F. Ahmed, “Predictors of Conversion to Multiple Sclerosis in Patients with Clinical Isolated Syndrome Using the 2010 Revised McDonald Criteria,” ISRN Neurology, vol. 2012, Article ID 792192, 6 pages, 2012. doi:10.5402/2012/792192

23. Dobson R, Ramagopalan S, Giovannoni G. The effect of gender in clinically isolated syndrome (CIS): a meta-analysis. Mult Scler 2012;18: 600–4. doi: 10.1177/1352458511426740.

24. Koch-Henriksen N, Sorensen PS. The changing demographic pattern of multiple sclerosis epidemiology. Lancet Neurol 2010; 9: 520–32. doi: 10.1016/S1474-4422(10)70064-8.

25. Mar Tintore, alex Rovira, Jordi Rio, Susana Otero-Romero, Georgina Arrambide, Carmen Tur et. al. Defining high, medium and low impact prognostic factors for developing multiple sclerosis. Brain 2015: 138; 1863–1874. doi:10.1093/brain/awv105

